# Unconventional kinetochore kinases KKT10/19 promote the metaphase to anaphase transition in *Trypanosoma brucei*

**DOI:** 10.1101/806224

**Authors:** Midori Ishii, Bungo Akiyoshi

## Abstract

The kinetochore is a macromolecular protein complex that drives chromosome segregation in eukaryotes. Unlike most eukaryotes that have canonical kinetochore proteins, evolutionarily divergent kinetoplastids such as *Trypanosoma brucei* have unconventional kinetochore proteins. *T. brucei* also lacks a canonical spindle checkpoint system and it therefore remains unknown how mitotic progression is regulated in this organism. Here we characterized two paralogous kinetochore proteins with a CLK-like kinase domain, KKT10 and KKT19, which localize at kinetochores in metaphase but disappear at the onset of anaphase. We found that these proteins are functionally redundant. Double knockdown of KKT10/19 led to a significant delay in the metaphase to anaphase transition. A kinase-dead mutant of KKT10 failed to rescue the KKT10/19 depletion phenotype, suggesting that its kinase activity is essential. We also found that phosphorylation of two kinetochore proteins KKT4 and KKT7 depends on KKT10/19 in vivo. Finally, we showed that the N-terminal part of KKT7 directly interacts with KKT10 and that kinetochore localization of KKT10 depends not only on KKT7 but also on the KKT8 complex. Our results reveal that kinetochore localization of KKT10/19 is tightly controlled to regulate the metaphase to anaphase transition in *T. brucei*.

**Summary:** *Trypanosoma brucei* has unique kinetochore proteins and lacks a canonical spindle checkpoint system. How mitotic progression is regulated in this organism remains unclear. Here we show that two redundant protein kinases KKT10/19 promote the metaphase to anaphase transition.

## Introduction

Proper segregation of chromosomes into two daughter cells during cell division is essential for the survival of all eukaryotes. Chromosomes are replicated during S phase and linked together by cohesin complexes (Nasmyth and Haering, 2009). The kinetochore is a macromolecular protein complex, which attaches to the centromeric region of each chromosome and interacts with spindle microtubules (Santaguida and Musacchio, 2009; Cheeseman, 2014). Kinetochore-microtubule attachments are monitored by a feedback mechanism called the spindle checkpoint, which delays the metaphase to anaphase transition until all chromosomes are attached to spindle microtubules emanating from opposite poles (Hoyt et al., 1991; Li and Murray, 1991). Spindle checkpoint components, such as Mad1, Mad2, Mad3/BubR1, Bub1, Bub3, and Mps1 are recruited to unattached kinetochores, creating a signal that inhibits Cdc20, an activator of the ubiquitin ligase called the anaphase promoting complex/cyclosome (APC/C) (Musacchio and Salmon, 2007). Once all chromosomes are properly bi-oriented, the spindle checkpoint is satisfied and APC/C gets activated, which leads to the degradation of securin and cyclin B (Yamano, 2019). Degradation of securin leads to the cleavage of cohesin complexes and separation of sister chromatids (Yanagida, 2000; Nasmyth et al., 2000), while that of cyclin B promotes mitotic exit (Hershko, 1999).

Kinetochores in many eukaryotes consist of more than 40 different proteins, some of which are conserved even in diverse eukaryotes (Meraldi et al., 2006; van Hooff et al., 2017). However, none of canonical kinetochore proteins is found in a group of evolutionarily-divergent eukaryotes called kinetoplastids (Lowell and Cross, 2004; Berriman et al., 2005). In *T. brucei*, which is a kinetoplastid parasite that causes human African trypanosomiasis (sleeping sickness) in sub-Saharan Africa, a number of unique kinetochore proteins have been identified, including KKT1–20, KKT22–25, and KKIP1–12 (Akiyoshi and Gull, 2014; Nerusheva and Akiyoshi, 2016; D’Archivio and Wickstead, 2017; Brusini et al., 2019; Nerusheva et al., 2019). Furthermore, homologs of spindle checkpoint components are apparently absent in *T. brucei*, except for a Mad2-like protein. However, this protein localizes only at basal bodies, and Cdc20 does not have a well-conserved Mad2-binding motif (Akiyoshi and Gull, 2013). Consistent with these findings, depolymerization of spindle microtubules does not delay the metaphase to anaphase transition (Ploubidou et al., 1999), suggesting that *T. brucei* indeed lacks a canonical spindle checkpoint system. In contrast, *T. brucei* has functional homologs of other mitotic machineries such as the CDK/cyclin system (Hammarton et al., 2003; Tu and Wang, 2004) and the anaphase promoting complex/cyclosome (APC/C) (Kumar and Wang, 2005). It remains unclear whether there is any regulatory mechanism for mitotic progression in *T. brucei*.

Protein kinases are known to play regulatory roles at various cellular locations, including kinetochores. Among known kinetoplastid kinetochore proteins, four proteins have a kinase domain, namely KKT2, KKT3, KKT10, and KKT19 (Akiyoshi and Gull, 2014). Because these proteins are not present in humans, they are attractive drug targets against kinetoplastid parasites. Previous studies identified small molecules that inhibit KKT10 and KKT19 (Nishino et al., 2013; Saldivia et al., 2019; Torrie et al., 2019). KKT10 and KKT19 (also known as TbCLK1 and TbCLK2) are paralogous proteins, apparently made by a recent gene duplication event (Akiyoshi and Gull, 2014). Their kinase domain is 100% identical and is classified as a member of the cdc2-like (CLK) kinase subfamily (Parsons et al., 2005). Although inhibition of KKT10/19 by RNAi-mediated knockdown or chemical compounds severely affected cell growth (Alsford et al., 2011; Nishino et al., 2013; Akiyoshi and Gull, 2014; Jones et al., 2014; Saldivia et al., 2019), little is known about their molecular functions.

Here, we have characterized KKT10 and KKT19 in *T. brucei*. We show that they are functionally redundant in procyclic form cells. Their double knockdown causes a delay in the metaphase to anaphase transition without affecting the localization of other kinetochore proteins. We also show that KKT4 and KKT7 are phosphorylated in a KKT10-dependent manner. Furthermore, we identify KKT7 as a direct interaction partner of KKT10. Yet kinetochore localization of KKT10 depends not only on KKT7 but also on the KKT8 complex. Taken together, our data have revealed that KKT10 and KKT19 play essential regulatory roles in trypanosomes.

## Results

### KKT10 and KKT19 are functionally redundant

Previous studies showed that simultaneous knockdown of both KKT10 and KKT19 caused growth defects in procyclic (Akiyoshi and Gull, 2014) and bloodstream form cells (Nishino et al., 2013; Jones et al., 2014; Saldivia et al., 2019). Although KKT10 and KKT19 have an identical protein kinase domain (Fig. 1A), it remained unclear whether they have distinct functions. Previous studies showed that KKT10 was essential for the proliferation of bloodstream form cells, while KKT19 was not (Nishino et al., 2013; Jones et al., 2014; Saldivia et al., 2019). To investigate their functions in procyclic cells, we performed RNAi using a construct that specifically targeted KKT10. We first confirmed reduction of the YFP-KKT10 signal upon induction of RNAi (Fig. S1A). However, to our surprise, these KKT10-depleted cells grew normally (Fig. S1B). We obtained a similar result for KKT19 depletion (data not shown), suggesting that KKT10 and KKT19 may be functionally redundant in procyclic cells. To test this possibility, we made strains that lack KKT10 or KKT19. Both alleles of KKT10 or KKT19 coding regions were replaced with drug resistant gene cassettes by a PCR-based method (Merritt and Stuart, 2013) to make kkt10∆/kkt10∆ (kkt10∆) or kkt19∆/kkt19∆ (kkt19∆) cells. Consistent with our RNAi results, both kkt10∆ and kkt19∆ cells were viable, confirming that KKT10 and KKT19 are functionally redundant in procyclic cells. However, kkt19∆ cells grew slower compared to wild-type or kkt10∆ cells (Fig. 1B). We found that KKT19 proteins are more abundant than KKT10 in wild-type cells (Fig. 1C, Fig. S1C). The mild growth defect of kkt19∆ cells may therefore imply that the amount of KKT10 proteins was insufficient to ensure normal cell growth.

**Figure 1.**
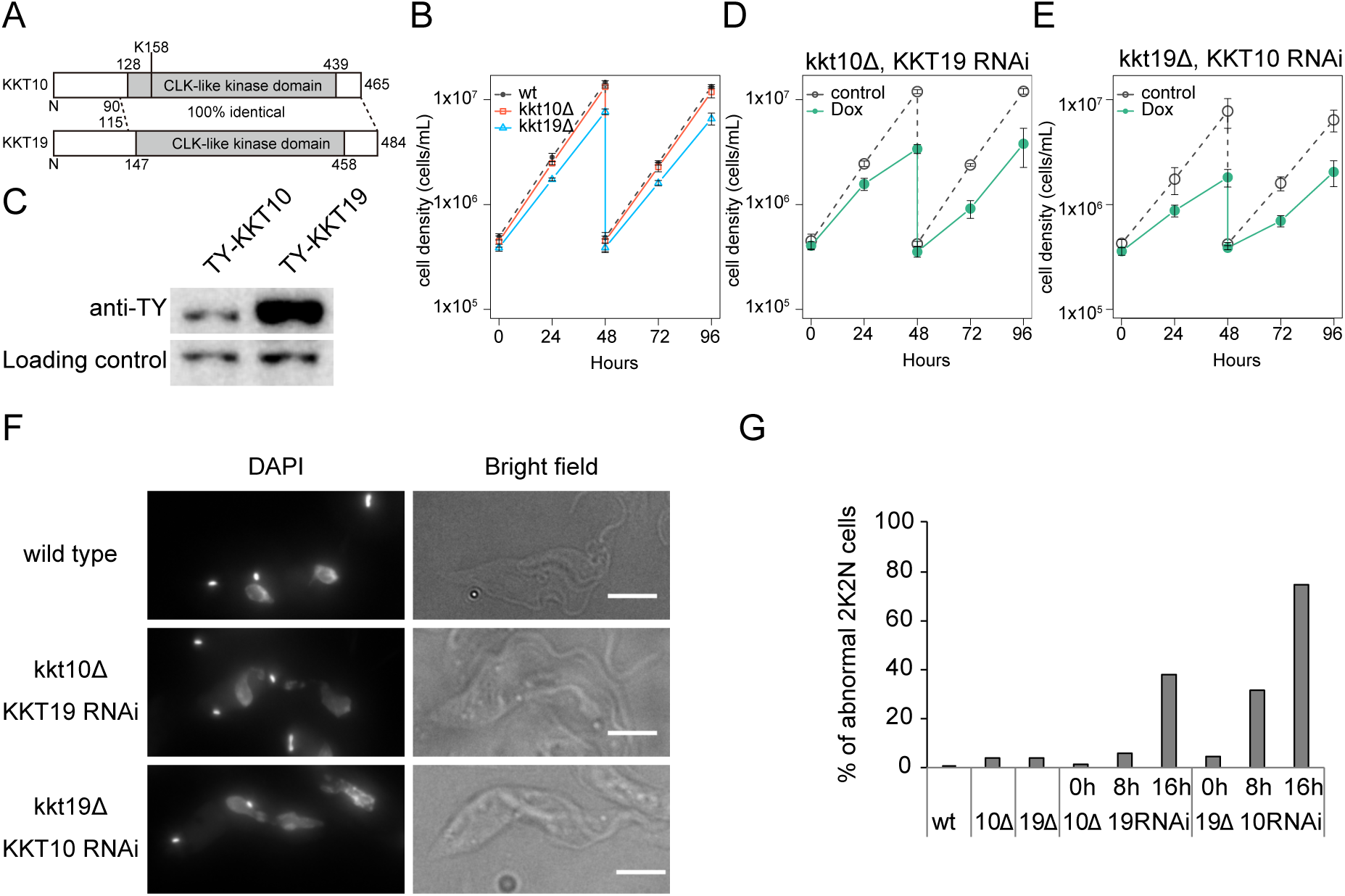
KKT10 and KKT19 are redundant. (A) Schematic representation of KKT10 and KKT19 proteins. (B) KKT10 (red) and KKT19 (blue) knockout cells are viable. Grey dashed line indicates a wild-type control. Error bars represent standard deviation from three independent experiments. Similar results were obtained from at least two independent clones. (C) Protein levels of TY-YFP-tagged KKT10 and KKT19 were monitored by immunoblotting. Representative of three independent experiments is shown. PFR2 was used as a loading control. Uncropped images are shown in Fig. S1. (D and E) RNAi-mediated knockdown of (D) KKT19 in kkt10 deletion cells and (E) KKT10 in kkt19 deletion cells affects cell growth. Control is an uninduced cell culture. Error bars represent standard deviation from three independent experiments. (F and G) KKT10/19 double depletion causes chromosome missegregation. (F) Examples of anaphase cells fixed at 16 hours postinduction of RNAi and stained with DAPI. Maximum intensity projections are shown. Bars, 5 µm. (G) Quantification of 2K2N cells with abnormal nuclear DNA morphology (n ≥ 74).

To deplete both KKT10 and KKT19 proteins, we performed KKT19-specific RNAi in kkt10∆ cells (Fig. 1D) or KKT10-specific RNAi in kkt19∆ cells (Fig. 1E), and found severe growth defects in both cases. To investigate the phenotype of KKT10 and KKT19 depletion, we examined chromosome segregation in anaphase. As we previously showed in KKT10/19 double knockdown cells (Akiyoshi and Gull, 2014), kkt10∆ KKT19 RNAi and kkt19∆ KKT10 RNAi cells had abnormal nuclear morphology in anaphase cells (Fig. 1F, G). In contrast, kkt10∆ or kkt19∆ cells had only small numbers of mis-segregation (Fig. 1G). These results confirm that KKT10 and KKT19 are functionally redundant in procyclic cells.

### Kinase activity of KKT10 is essential for cell proliferation

To investigate the importance of KKT10/19 kinase activities, we made a kinase-dead form of KKT10 by mutating lysine 158 (Fig. 1A), a residue conserved in active protein kinases in eukaryotes, in the N-terminal YFP-tagging construct for KKT10. We first confirmed that YFP-KKT10^WT^ was able to rescue the KKT10/19 double depletion phenotype (Fig. S2A). We then made a strain that has YFP-KKT10^K158A^ as the sole source of KKT10 in cells. YFP-KKT10^K158A^ localized normally at kinetochores from S phase to metaphase (Fig. 2A). However, we observed a severe growth defect upon depletion of KKT19 (Fig. 2B), which was almost comparable to the double knockdown of KKT10/19 (Fig. 1D, E). As expected, abnormal chromosome segregation was observed in KKT10 kinase-dead cells (Fig. 2C). These results show that the kinase activity is a major function of KKT10 but is not important for its own kinetochore localization.

**Figure 2.**
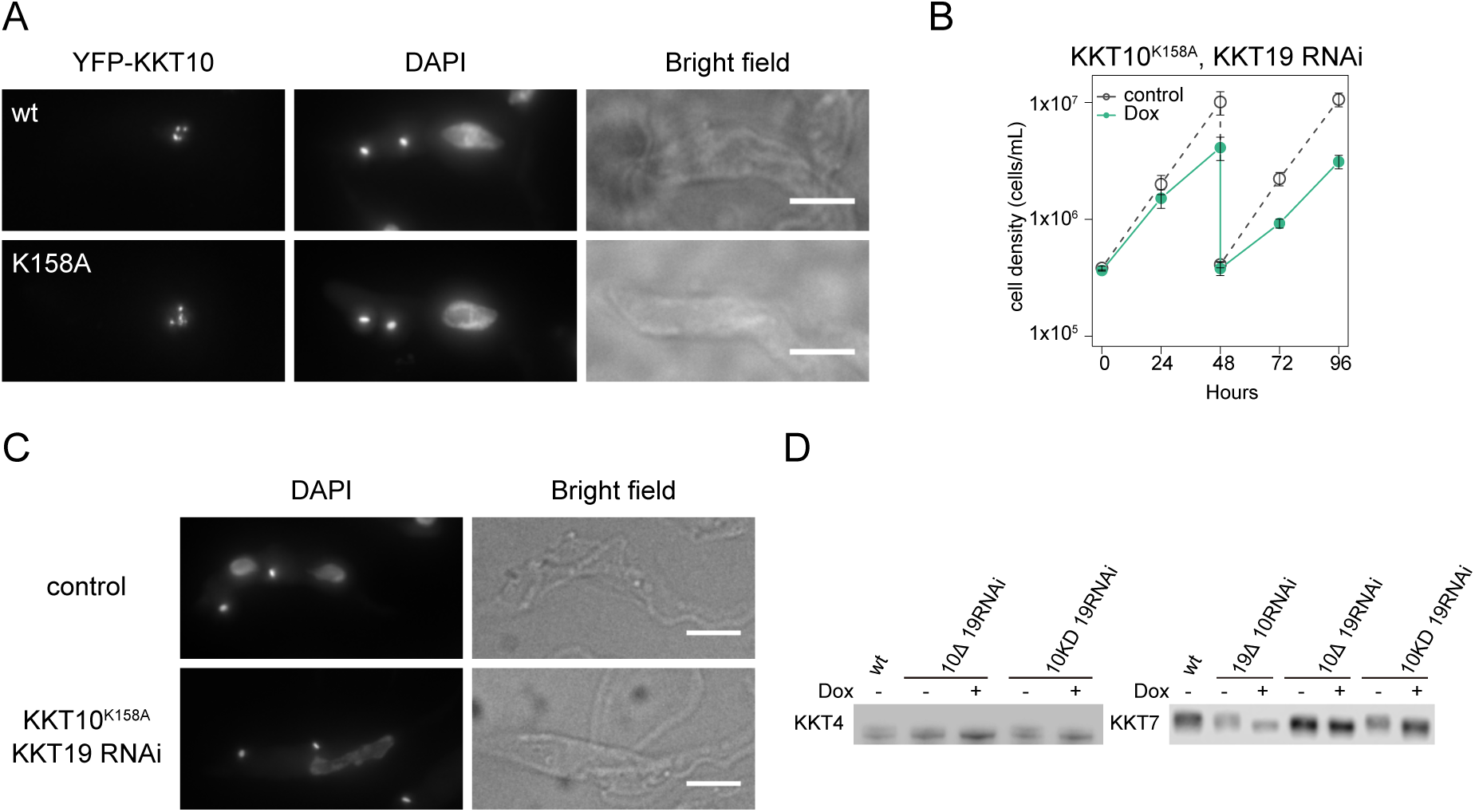
Phosphorylation of KKT4 and KKT7 depends on KKT10 kinase activities. (A) Examples of cells expressing YFP-KKT10 (wild type or K158A mutant) stained with DAPI. Maximum intensity projections are shown. Bars, 5 µm. (B) Expression of YFP-KKT10^K158A^ fails to rescue the KKT19 RNAi phenotype. Control is an uninduced cell culture. Error bars represent standard deviation from three independent experiments. Similar results were obtained from three clones. (C) Cells expressing YFP-KKT10^K158A^ in the kkt10∆ KKT19 RNAi strain were fixed at 24 hours postinduction of RNAi and stained with DAPI. Control is an uninduced cell culture. Maximum intensity projections are shown. Bars, 5 µm. (D) Phosphorylation of KKT4 and KKT7 depends on KKT10/19. 3Flag-tagged KKT4 and KKT7 were detected upon induction of RNAi for 24 hours. Uncropped images are shown in Fig. S2. Representative of at least three independent experiments is shown.

To identify the substrate of KKT10/19 kinases, we searched protein sequences of kinetochore proteins that have consensus phosphorylation sequences of KKT10/19 (RxxS) (Torrie et al., 2019). KKT4 and KKT7 have several such motifs, so we tested whether they are phosphorylated in vivo by performing immunoblots against these proteins. We detected electrophoretic mobility shifts in wild-type cells, which disappeared in KKT10/19-depleted cells and KKT10 kinase-dead cells (Fig. 2D, Fig. S2B, C). These results show that KKT4 and KKT7 are phosphorylated in a KKT10/19-dependent manner in vivo.

### Metaphase to anaphase transition is delayed without KKT10/19

KKT10/19 localize at kinetochores from S phase until anaphase onset. This localization pattern is reminiscent of that of spindle checkpoint proteins in other eukaryotes despite the fact that *T. brucei* does not appear to have a functional spindle checkpoint (Ploubidou et al., 1999; Hayashi and Akiyoshi, 2018). Although it has been shown that inhibition of KKT10/19 results in cell cycle defects in bloodstream form cells (Jones et al., 2014; Saldivia et al., 2019), it remains unclear whether KKT10/19 have a direct role in cell cycle regulation. To address this question, we examined the cell cycle status of KKT10/19 knockdown cells (Fig. S3A) (Akiyoshi and Gull, 2014). *T. brucei* has a characteristic DNA structure called the kinetoplast which contains mitochondrial DNA. Kinetoplasts segregate prior to the nuclear division, thus the number of kinetoplasts (K) and nuclei (N) in a cell indicates the cell cycle stage: 1K1N (one kinetoplast and one nucleus) for G1 to S phase, 2K1N (two kinetoplasts and one nucleus) for G2 to metaphase, and 2K2N (two kinetoplasts and two nuclei) for anaphase to telophase (Robinson et al., 1995). We found that the ratio of 1K1N cells decreased, while that of 2K1N cells increased in KKT10/19 knockdown cells at 24 hours post-induction (Fig. 3A). We also analyzed the cell cycle profile in kkt10∆ KKT19 RNAi and kkt19∆ KKT10 RNAi cells, and obtained similar results (Fig. S3B, C). These results suggest that there is a delay in nuclear division upon depletion of KKT10/19. To directly test this possibility, we monitored YFP-tagged cyclin B (CYC6) that appears in the nucleus in G2 and disappears at the onset of anaphase (Hayashi and Akiyoshi, 2018). We found that the number of CYC6-positive 2K1N cells increased in KKT10/19 knockdown cells (Fig. 3B), confirming the delay in the metaphase to anaphase transition.

**Figure 3.**
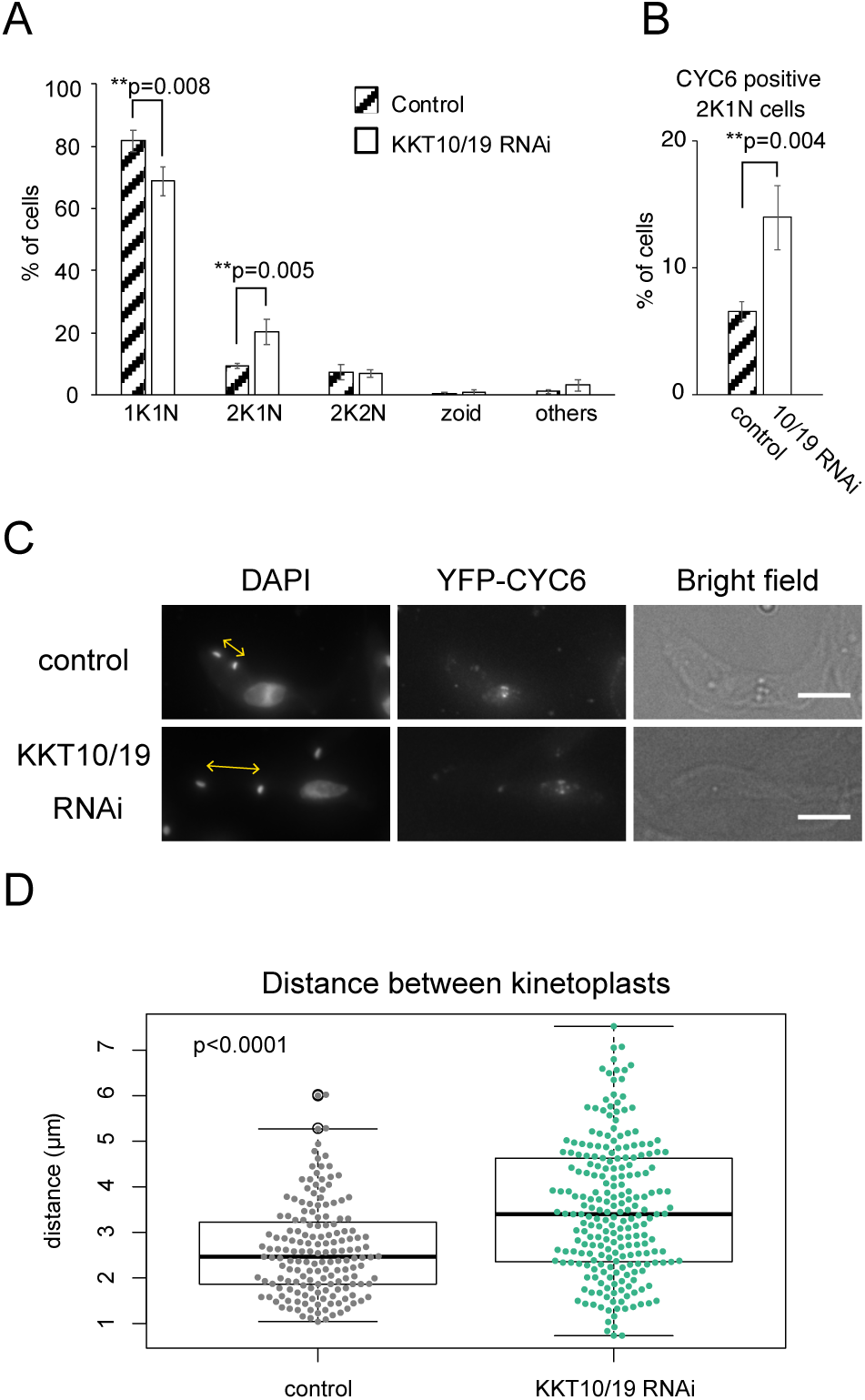
KKT10/19 depletion delays the metaphase to anaphase transition. (A and B) Quantification of (A) cells with indicated DNA contents, or (B) 2K1N cells that have nuclear CYC6 signals. Cells were fixed at 24 hours postinduction of KKT10/19 RNAi. Control is an uninduced cell culture. P-values were calculated by Student’s t-test, one-tailed distribution. Error bars represent standard deviation from three independent experiments (n ≥ 246). (C) KKT10/19-depleted cells have longer distance between kinetoplastids. Cells were fixed at 24 hours postinduction of RNAi and stained with DAPI. Control is an uninduced cell culture. Maximum intensity projections are shown. Bars, 5 µm. (D) Quantification of distance between two kinetoplasts in nuclear CYC6-positive 2K1N cells upon KKT10/19 RNAi. Data were collected from cells at 24 hours postinduction. Control is an uninduced cell culture. p<0.0001 was calculated by Welch two sample t-test (n ≥ 167).

In *T. brucei*, the distance between two kinetoplastids is another cell cycle marker, which gets longer as cell cycle progresses and becomes maximum before cytokinesis (Robinson et al., 1995). Although nuclear and cytoplasmic events are coordinated under proliferating conditions, inhibition of nuclear division does not prevent the progression of cytoplasmic events (Ploubidou et al., 1999). We therefore measured the distance between the two kinetoplasts in 2K1N (G2/M) cells that have positive CYC6 signals to examine their cytoplasmic cell cycle status (Fig. 3C, D). The average distance between the two kinetoplasts was 2.7 µm in control cells compared to 3.5 µm in KKT10/19 knockdown cells (Fig. 3D). This result further supports our finding that the metaphase to anaphase transition in the nucleus is delayed in KKT10/19 knockdown cells.

### KKT10/19 are dispensable for the localization of other kinetochore proteins

It was recently shown that treatment of bloodstream form cells with AB1, a covalent kinase inhibitor against KKT10, affected the localization of some kinetochore proteins (Saldivia et al., 2019). Given its potential off-target effects, however, it remains unclear whether the observed defects were actually due to inhibition of KKT10/19. To test whether KKT10/19 regulate the localization of kinetochore proteins in procyclic cells, YFP-tagged KKT1, KKT4, KKT7, KKT8, KKT14 and KKIP1 were imaged in kkt10LΔ KKT19 RNAi cells (Fig. 4). These proteins localize at kinetochores at different cell cycle stages in wild-type cells. KKT4, a microtubule-binding component, localizes at kinetochores constitutively (Akiyoshi and Gull, 2014; Llauró et al., 2018). KKT1, KKT7, and KKIP1 localize at kinetochores from S phase to anaphase, while KKT8 localizes from S to metaphase (Akiyoshi and Gull, 2014; D’Archivio and Wickstead, 2017). KKT14 localizes from G2 to anaphase (Akiyoshi and Gull, 2014). We found that although severe chromosome segregation defects were observed, kinetochore localization of these proteins was not perturbed by KKT10/19 knockdown (Fig. 4). Therefore, KKT10/19 are not essential for the localization of kinetochore proteins in procyclic cells.

**Figure 4.**
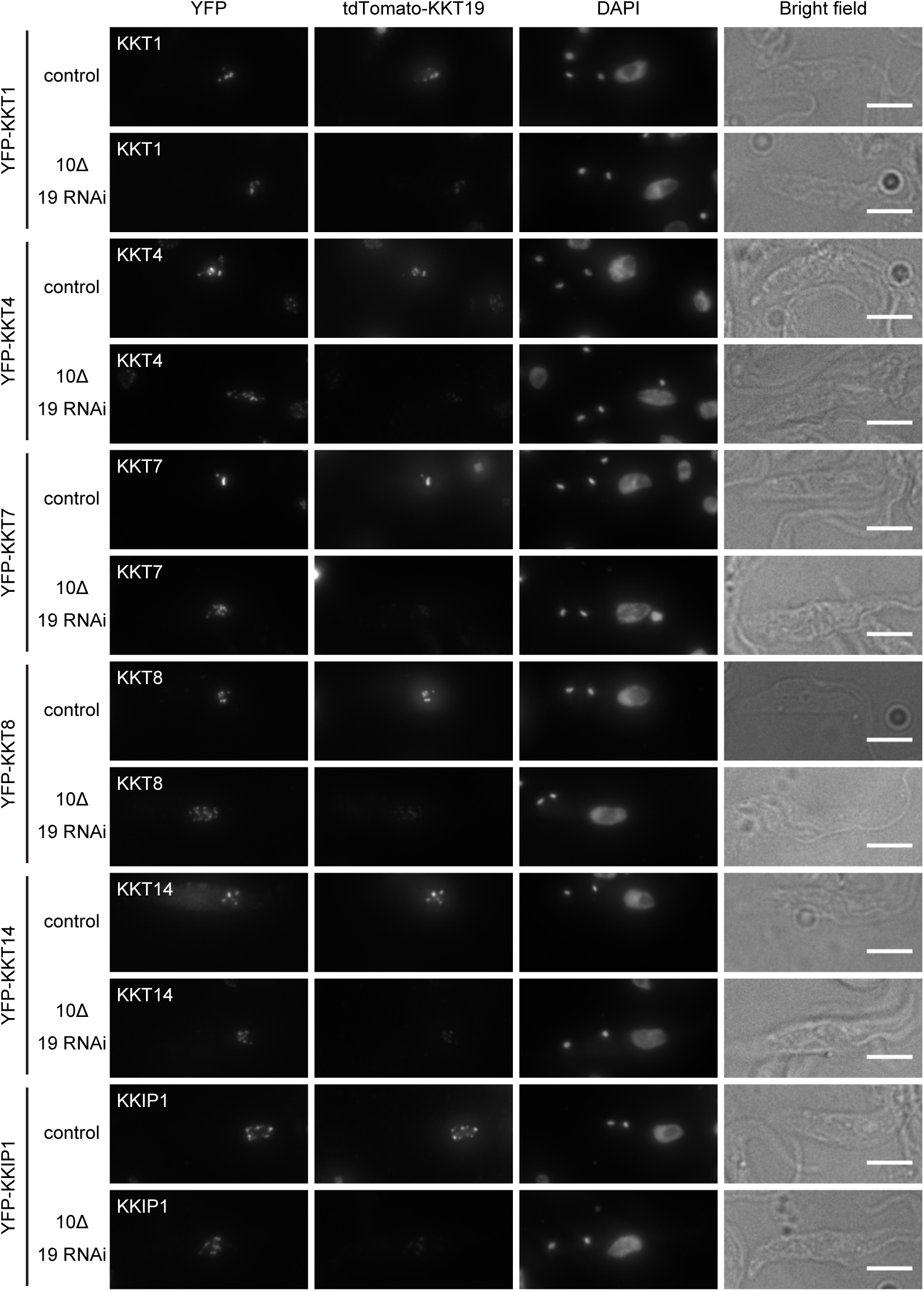
KKT10/19 are dispensable for the localization of other kinetochore proteins. YFP-tagged KKT1, 4, 7, 8, 14 or KKIP1 were imaged after KKT10/19 depletion. Examples of 2K1N cells fixed at 24 hours postinduction of RNAi and stained with DAPI are shown. Control is an uninduced cell culture. Maximum intensity projections are shown. Bars, 5 µm.

### N-terminal part of KKT7 recruits KKT10/19 onto kinetochores

We next aimed to reveal how KKT10/19 are recruited to kinetochores and how their localization is regulated. Our previous data showed that KKT7 was one of the most abundant proteins that co-purified with YFP-tagged KKT10 or KKT19 (Akiyoshi and Gull, 2014). During the course of our studies on KKT7, we found that its N-terminal region (residues 2–261: KKT7N) or the C-terminal region (262–644: KKT7C) can localize at kinetochores when ectopically expressed in wild-type trypanosomes (Fig. 5A, B). Immunoprecipitation of these fragments revealed that KKT7N co-purified with KKT10 and KKT19, while KKT7C co-purified with a number of kinetochore proteins, but not with KKT10 or KKT19 (Fig. 5C, D). These results indicate that KKT7 is recruited to kinetochores via its C-terminal part and that KKT10 is recruited by the N-terminal part of KKT7. Kinetochore localization of KKT7N in metaphase, but not in anaphase when KKT10/19 disappear from kinetochores, supports this idea. To test whether the KKT7 N-terminal region is sufficient to recruit KKT10, we used a LacO-LacI system (Landeira and Navarro, 2007) to tether KKT7N to an ectopic locus. We found that KKT10 and KKT19 are recruited to the KKT7N-LacI protein on the LacO locus (Fig. 5E). We then tested whether KKT10 directly interacts with KKT7N. We co-expressed 6HIS-KKT10 and KKT7N in *E. coli* and performed metal affinity chromatography, revealing that KKT7N co-purifies with 6HIS-KKT10 (Fig. 5F). Finally, we tested whether the localization of KKT10 depends on KKT7. We found that, in KKT7-depleted cells, the YFP-KKT10 signal was mostly diffuse in the nucleus, occasionally with few dots (Fig. 5G). Taken together, these results establish that KKT10/19 are recruited to the kinetochore by interacting with the N-terminal region of KKT7.

**Figure 5.**
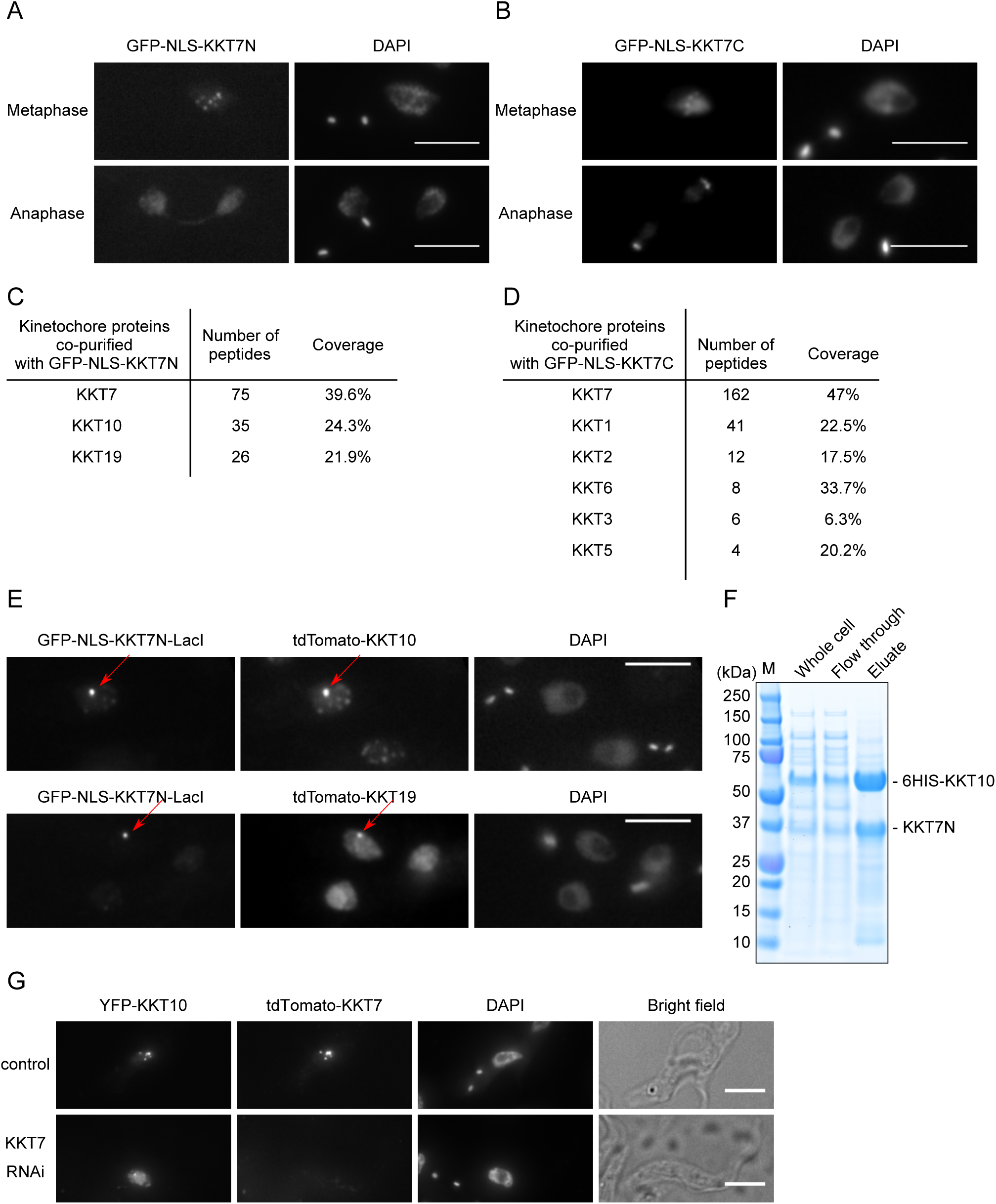
KKT7 binds and recruits KKT10/19 onto kinetochores. (A) Ectopically expressed GFP-NLS-KKT7N^2–261^ localizes at kinetochores in metaphase, not in anaphase. (B) Ectopically expressed GFP-NLS-KKT7C^262–644^ localizes at kinetochores in metaphase and anaphase. (C) KKT7N^2–261^ co-purifies with KKT10 and KKT19, but not with other kinetochore proteins. See Supplementary Table S2 for all proteins identified by mass spectrometry. (D) KKT7C^262–644^ co-purifies with several kinetochore proteins, but not with KKT10/19. (E) KKT7N^2–261^ is sufficient to recruit KKT10 and KKT19 to a non-centromeric locus in vivo. For (A–E), inducible GFP fusion proteins were expressed with 10 ng/mL doxycycline for 24 hours. Scale bars, 5 µm. (F) KKT7N^2–261^ directly interacts with KKT10. Recombinant KKT7N^2–261^ and 6HIS-KKT10 proteins were co-expressed in *E. coli*, followed by metal affinity chromatography. (G) Localization of YFP-tagged KKT10 is affected in KKT7 knockdown cells. Similar results were obtained in 88% of 2K1N cells (n = 26). Cells were fixed at 24 hours postinduction of RNAi and stained with DAPI. Control is an uninduced cell culture. Maximum intensity projections are shown. Bars, 5 µm.

### Localization of KKT10 is also dependent on KKT9 and KKT11

Regulation of spatio-temporal localization of a protein is often linked with its functional regulation (Dou et al., 2019). KKT10 and KKT19 localize at kinetochores from S phase until the onset of anaphase. This localization pattern is also found in four other kinetochore proteins, KKT8, KKT9, KKT11, and KKT12 (Akiyoshi and Gull, 2014). We used a bacterial co-expression system and found that KKT9/11/12 co-purified with 6HIS-KKT8, suggesting that they form a complex (Fig. 6A). We call this four-protein complex, the KKT8 complex. Given its similar localization pattern with KKT10/19, we next tested whether the KKT8 complex regulates the localization of KKT10 by performing RNAi against two of its components, KKT9 and KKT11, for which efficient depletions were achieved. We found that YFP-KKT10 failed to localize at kinetochores properly in both cases (Fig. 6B). Therefore, kinetochore localization of KKT10 relies not only on KKT7 but also the KKT8 complex.

**Figure 6.**
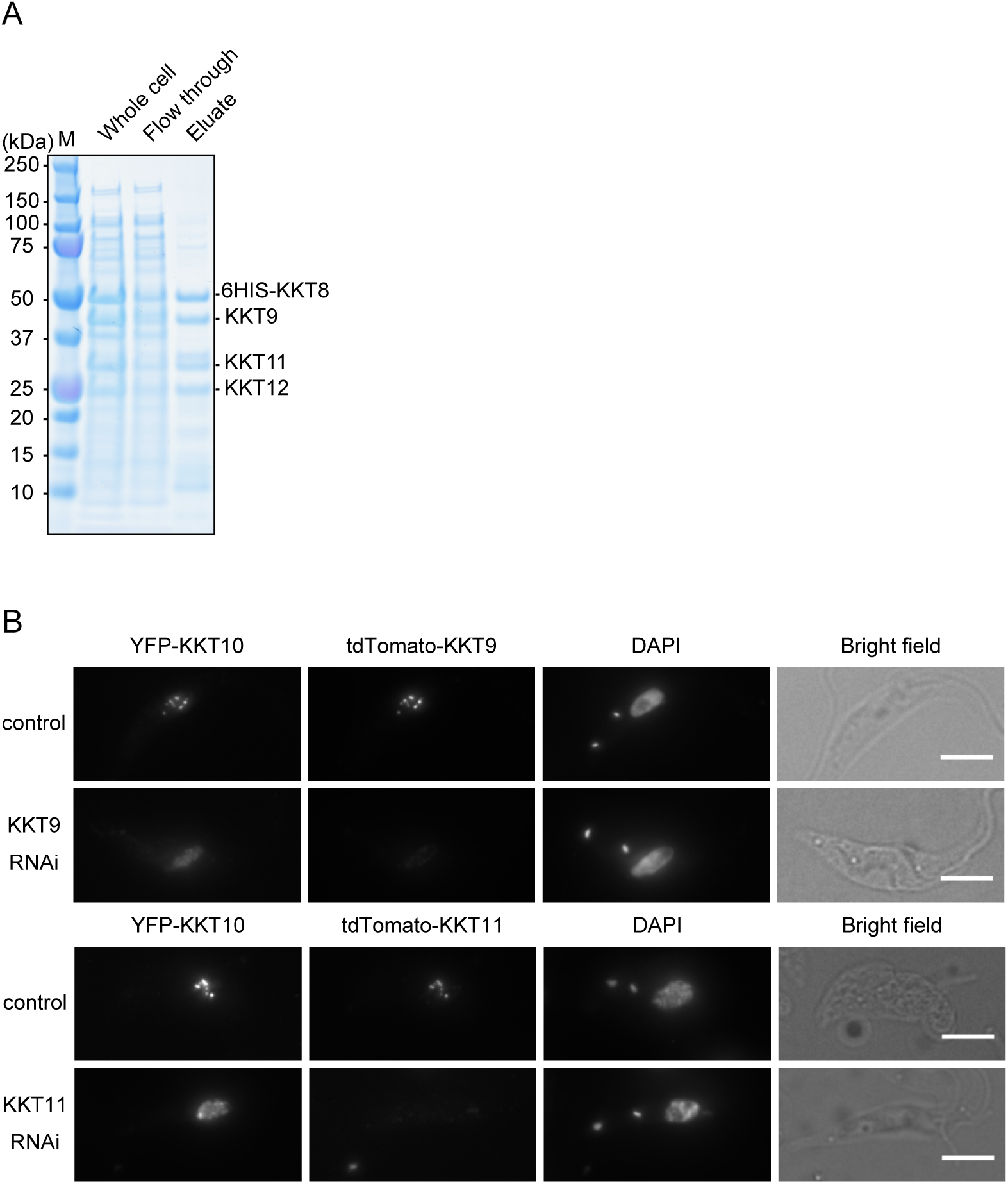
KKT10 localization is also dependent on the KKT8 complex. (A) KKT8, KKT9, KKT11, and KKT12 form a complex. Recombinant 6HIS-KKT8, KKT9, KKT11, and KKT12 proteins were co-expressed in *E. coli*, followed by metal affinity chromatography. (B) Localization of YFP-tagged KKT10 is affected in KKT9 and KKT11 knockdown cells. Similar results were obtained in more than 75% of 2K1N cells (n ≥ 16). Cells were fixed at 24 hours postinduction of RNAi and stained with DAPI. Control is an uninduced cell culture. Maximum intensity projections are shown. Bars, 5 µm.

## Discussion

Protein kinases play three major regulatory roles at kinetochores in eukaryotes: kinetochore assembly (e.g. CDK, Aurora B), error correction (e.g. Aurora B, Mps1), and the spindle checkpoint (e.g. Mps1, Bub1) (London and Biggins, 2014; Musacchio and Desai, 2017). While apparent homologs of Mps1 or Bub1 are absent, CDK (Hammarton et al., 2003; Tu and Wang, 2004) and Aurora B (Li et al., 2008) are present in *T. brucei*. In addition, there are four protein kinases that specifically localize at kinetochores. KKT2 and KKT3, whose kinase domains are classified as unique among eukaryotic kinase subfamilies, are putative DNA-binding kinetochore proteins that localize at kinetochores throughout the cell cycle. In contrast, KKT10 and KKT19, which have a CLK-like kinase domain, localize at kinetochores from S phase until the onset of anaphase. In other eukaryotes, CLK kinases are implicated in RNA splicing (Corkery et al., 2015), not kinetochore functions, suggesting that KKT10/19 likely evolved from a CLK kinase to carry out distinct functions in kinetoplastids.

*T. brucei* has a complex life cycle to survive in both animal hosts and insect vectors, and cells change many biological features to adapt to different environments (Matthews, 2005). Our findings in procyclic cells show that KKT10 and KKT19 are functionally redundant and that KKT19 is more abundant than KKT10. In contrast, previous studies in bloodstream form cells showed that KKT10 was essential for proliferation, while KKT19 was not, raising a possibility that KKT10 may be more abundant than KKT19 in this life stage. Alternatively, it is possible that KKT10 and KKT19 have non-overlapping functions in bloodstream form cells.

Saldivia et al. recently found that a covalent kinase inhibitor AB1 (Lelais et al., 2016) has potent activity against several trypanosomatids and identified KKT10 as a target in bloodstream form *T. brucei* (Saldivia et al., 2019). They also found that KKT10 phosphorylated KKT2 at serine 507 and/or serine 508 and that the KKT2^S507A^ ^S508A^ phospho-deficient mutant failed to rescue the growth defect caused by KKT2 RNAi in bloodstream form cells. They further showed the phosphorylation at these residues by KKT10 was important for recruiting some kinetochore proteins to kinetochores (Saldivia et al., 2019). However, we did not find significant localization defects of kinetochore proteins in KKT10/19-depleted procyclic cells. Moreover, the KKT2^S507A^ ^S508A^ mutant localized normally at kinetochores and supported cell growth upon induction of KKT2 RNAi (Fig. S4). We do not know the underlying molecular mechanisms for the observed differences between bloodstream and procyclic form cells. Further study is required to understand stage-specific and common mechanisms of kinetochore assembly in *T. brucei*.

In other eukaryotes, the spindle checkpoint monitors attachment errors and delays mitotic progression by inhibiting Cdc20, an activator of APC/C. *T. brucei* lacks canonical spindle checkpoint components but has APC/C. Building on previous studies that noted cell cycle defects upon KKT10 knockdown in bloodstream form cells (Jones et al., 2014; Saldivia et al., 2019), we used cyclin B as a nuclear cell cycle marker and showed that KKT10/19 are important for the metaphase to anaphase transition in procyclic cells. Interestingly, one of their targets, KKT4, has microtubule-binding activities (Llauró et al., 2018) and co-purifies with some APC/C subunits (Akiyoshi and Gull, 2014). We speculate that phosphorylation of KKT4 may change depending on its microtubule attachment status, directly affecting the interaction between KKT4 and APC/C subunits. The molecular mechanism of this potentially new form of mitotic regulation will need to be investigated in the future for better understanding of cell cycle progression in *T. brucei*.

## Acknowledgment

We thank Miguel Navarro for providing Mig75 and Mig96 plasmids, Song Tan for the pST44 plasmid, Keith Gull for BB2 and L8C4 antibodies, and Sam Dean for pPOT plasmids. We also thank the Advanced Proteomics Facility for mass spectrometry analysis, as well as the Micron Oxford. Midori Ishii was supported by a long-term fellowship from the TOYOBO Biotechnology Foundation in 2017. Bungo Akiyoshi was supported by a Wellcome Trust Senior Research Fellowship (grant no. 210622/Z/18/Z) and the European Molecular Biology Organization Young Investigator Program.

## Materials and methods

### Primers and plasmids

Primers, plasmids, and synthetic DNA fragments used in this study are listed in Supplementary Table S1. To make head-to-head RNAi constructs (pBA188; KKT19-specific and pBA190; KKT10-specific), gene fragments (KKT19 26–338 bp and KKT10 60–286 bp) were amplified with primers BA557/BA558 (KKT19) and BA561/BA562 (KKT10) from genomic DNA, and cloned into p2T7-177 (Wickstead et al., 2002) using *Spe*I/*Hin*dIII. To make hairpin RNAi constructs (pBA865; KKT7, pBA866; KKT10-specific, pBA1200; KKT11, and pBA1710; KKT2), synthetic DNA fragments (BAG28; KKT7 33–489 bp, BAG29; KKT10 14–286 bp, BAG61; KKT11 99–598 bp, and BAG94; KKT2 5’UTR 342 bp (GeneArt)) were cloned into pBA310 (Nerusheva and Akiyoshi, 2016) using *Hin*dIII/*Bam*HI. To make pBA1356 (TY-YFP-KKT10), the N-terminal region of the KKT10 coding sequence (4–1000 bp) was amplified with primers BA294/BA1874 from genomic DNA, and cloned into pBA74 (Akiyoshi and Gull, 2014) using *Xba*I/*Not*I. To make pBA1357 (TY-YFP-KKT10^K158A^), site-directed mutagenesis was performed using primers BA1875/BA1876 and pBA1356 as a template. To make N-terminal TY-tdTomato-tagging constructs (pBA1373; KKT19, pBA1419; KKT7, pBA1420; KKT8, pBA1586; KKT9 and pBA1587; KKT11), endogenous gene targeting sequences from TY-YFP-tagging constructs (pBA100; KKT19, pBA72; KKT7, pBA68; KKT8, pBA72; KKT9 and pBA75; KKT11 (Akiyoshi and Gull, 2014)) were subcloned into pBA148 using *Xba*I/*Bam*HI. To make pBA1444 (3Flag-tagging), PCR was performed using primers BA1995/BA1996 and pEnT6B-Y (Kelly et al., 2007) as a template, then the DNA was digested with *Xba*I and self-ligated. To make N-terminal 3Flag-tagging constructs (pBA1452; KKT4 and pBA1453; KKT7), endogenous gene targeting sequences from pBA71; KKT4 and pBA72; KKT7 (Akiyoshi and Gull, 2014) were subcloned into pBA1444 *Xba*I/*Bam*HI sites. To make pBA1806 (N-terminal TY-YFP-tagging construct for KKT2), the coding sequence of KKT2 (4–3780 bp) was amplified with primers BA266/BA2346 from genomic DNA and subcloned into *Hin*dIII/*Not*I sites of pBA67 (Akiyoshi and Gull, 2014), and then site-directed mutagenesis was performed using primers BA2639/BA2640 to make pBA2034. To make pBA607, 6HIS-KKT10 was amplified from a plasmid that encodes KKT10 using BA1061/BA1062 and cloned into pST44 (Tan et al., 2005) using *Xba*I/*Stu*I sites. To make pBA660, KKT7N (2–261) was amplified from a plasmid that encodes KKT7 using BA1065/BA1068 and cloned into pBA607 using *Eco*RI/*Sac*I sites. To make pBA457, synthetic DNA (GeneArt) that has four translation cassettes for full-length 6HIS-KKT8, KKT9, KKT11, and KKT12 (each gene was codon-optimized for expression in *E. coli*) was subcloned into pST44 using *Xba*I/*Mlu*I sites. To make pBA346, a DNA fragment for the KKT7 N-terminal domain was amplified using primers BA733/BA734 and cloned into pBA310 *Pac*I/*Afl*II sites. To make pBA347, a DNA fragment for the KKT7 C-terminal domain was amplified using primers BA735/BA736 and cloned into pBA310 *Pac*I/*Afl*II sites. An inducible GFP-NLS-LacI plasmid (pBA795) was made as follows. First, LacI was amplified from pMig75 (Navarro and Gull, 2001) using primers BA1063/BA1064 and cloned into pBA310 using *Asc*I/*Afl*II to make pBA608. Then the DNA fragment containing GFP-NLS-LacI was subcloned into pDex877-GFP-TY using *Nhe*I/*Not*I to make pBA795. DNA fragment containing the KKT7 N-terminal domain was amplified with primers BA1401/BA1402 and cloned into pBA795 at *Pac*I/*Asc*I site to make pBA891. To make pBA892 (N-terminal TY-tdTomato-tagging vector with hygromycin marker), DNA fragment of pBA148 that encodes TY-tdTomato was subcloned into pEnT5-Y using *Spe*I/*Xba*I sites. To make N-terminal TY-tdTomato-tagging constructs (pBA919; KKT10 and pBA1103; KKT19), endogenous gene targeting sequences from TY-YFP-tagging constructs (pBA74; KKT10 and pBA100; KKT19 (Akiyoshi and Gull, 2014)) were subcloned into pBA892 using *Xba*I/*Bam*HI. All constructs were sequence verified.

### Cells

All cell lines used in this study were derived from *T. brucei* SmOxP927 procyclic form cells (Poon et al., 2012). and listed in Supplementary Table S1. Cells were grown at 28°C in SDM-79 medium (Life Technologies) supplemented with 10% (v/v) heat-inactivated fetal calf serum (Sigma) (Burn and Schonenberger, 1979) with puromycin (Sigma) and appropriate drugs. For induction of RNAi or ectopic expression of GFP-NLS fusion proteins, doxycycline (Sigma) was added to the medium to a final concentration of 1 µg/mL or 10 ng/mL, respectively.

Gene deletions were carried out as described previously (Merritt and Stuart, 2013). To make deletion strains (BAP1004; kkt19∆, BAP1054; kkt10∆), fusion PCR of three PCR products (1. Upstream targeting sequence distal to KKT19 (primers BA1848/BA1849) or KKT10 (primers BA1836/BA1837) amplified from genomic DNA, 2. Neomycin marker cassette amplified from pBA183 using primers BA903/BA904, 3. Downstream targeting sequence distal to KKT19 (primers BA1855/BA1856) or KKT10 (primers BA1844/BA1845) amplified from genomic DNA) were transfected into SmOxP927 by electroporation. Transfected cells were selected by addition of 30 µg/mL G418 (Sigma) and cloned by dispensing dilutions into 96-well plates. To make BAP1068 (kkt19∆/kkt19∆) and BAP1073 (kkt10∆/kkt10∆), fusion PCR of three PCR products (1. Upstream targeting sequence distal to KKT19 (primers BA1858/BA1859) or KKT10 (primers BA1838/BA1839) amplified from genomic DNA, 2. clonNAT marker cassette amplified from pMig75 using primers BA905/BA906, 3. Downstream targeting sequence distal to KKT19 (primers BA1860/BA1861) or KKT10 (primers BA1842/BA1843) amplified from genomic DNA) were transfected into BAP1004 or BAP1054. Transfected cells were selected by addition of 100 µg/mL nourseothricin/clonNAT (Jena Bioscience) and cloned by dispensing dilutions into 96-well plates. Deletions were checked by PCR. We could not maintain the strain that has 3Flag-KKT4 kkt19Δ/kkt19Δ KKT10 RNAi because cells were quite sick, which is likely due to a small level of leakage of RNAi even in the absence of doxycycline (data not shown). This result suggests that there is a negative genetic interaction between 3FLAG-KKT4 and KKT10/19.

For C-terminally YFP-tagged KKT2 (BAP1579), YFP-tagging cassette was amplified from pPOTv7 (Dean et al., 2015) using primers BA2267/BA2268. The PCR product was transfected into SmOxP927 by electroporation. Transfected cells were selected by the addition of 10 µg/mL blasticidin S (Insight biotechnology).

All plasmids were linearized by *Not*I and transfected to trypanosomes by electroporation into an endogenous locus (TY-YFP tagging, TY-tdTomato tagging, 3Flag-6His-YFP tagging, and 3Flag tagging), rDNA locus (pMig96), or 177bp repeats on minichromosomes (RNAi, GFP-NLS-KKT7N/C, and GFP-NLS-KKT7N-LacI). Transfected cells were selected by the addition of 25 µg/mL hygromycin (Sigma), 10 µg/mL blasticidin S (Insight biotechnology) or 5 µg/mL phleomycin (Sigma).

### Fluorescence microscopy

Cells were fixed with formaldehyde as previously described (Nerusheva and Akiyoshi, 2016). Images were captured at room temperature on a DeltaVision fluorescence microscope (Applied Precision) installed with softWoRx version 5.5 housed in the Oxford Micron facility. Fluorescent images were captured with a CoolSNAP HQ camera using 60x objective lenses (1.42 NA). Typically, 25 optical slices spaced 0.2 µm apart were collected. Maximum intensity projection images were generated by Fiji software (Schindelin et al., 2012).

### Immunoblotting

Cells were harvested by centrifugation (800g, 5 min) and washed with 1 mL PBS. Then 25% trichloroacetic acid (TCA) was added and mixture was incubated for 5 min on ice, followed by centrifugation at 14,000 rpm at room temperature for 1 min. Then cells were washed with ice-cold acetone and the supernatant removed. The pellet was resuspended in 1xLDS sample buffer (Invitrogen) with 0.1 M DTT. Denaturation of proteins was performed for 5 min at 95°C.

SDS-PAGE and immunoblots were performed by standard methods using the following mouse monoclonal antibodies: BB2 (anti-TY, 1: 100) (Bastin et al., 1996) for TY-YFP-tagged KKT proteins, L8C4 (anti-PFR2, 1: 1500) (Kohl et al., 1999) for a loading control, and anti-Flag (Sigma, clone M2 F3165, 1: 500). Bands were visualized by horseradish-peroxidase-conjugated sheep anti-mouse IgG antibodies (GE Healthcare).

### Protein purification

For affinity-purification of GFP-NLS-KKT7N and KKT7C from trypanosomes, their expression was induced with 10 ng/ml doxycycline for 24 hours. Immunoprecipitation and mass spectrometry were performed essentially as described previously (Nerusheva et al., 2019) at the Advanced Proteomics Facility in University of Oxford using MASCOT (Matrix Science). Proteins identified with at least two peptides were considered and shown in Supplementary Table S2.

Purification of recombinant proteins (pBA660: 6HIS-KKT10 KKT7N, pBA457: 6HIS-KKT8, KKT9, KKT11, KKT12) from bacteria were performed as previously described (Llauró et al., 2018) except that proteins were expressed in Rosetta 2(DE3)pLys cells (Novagen).

## Supplemental materials

**Figure S1.**
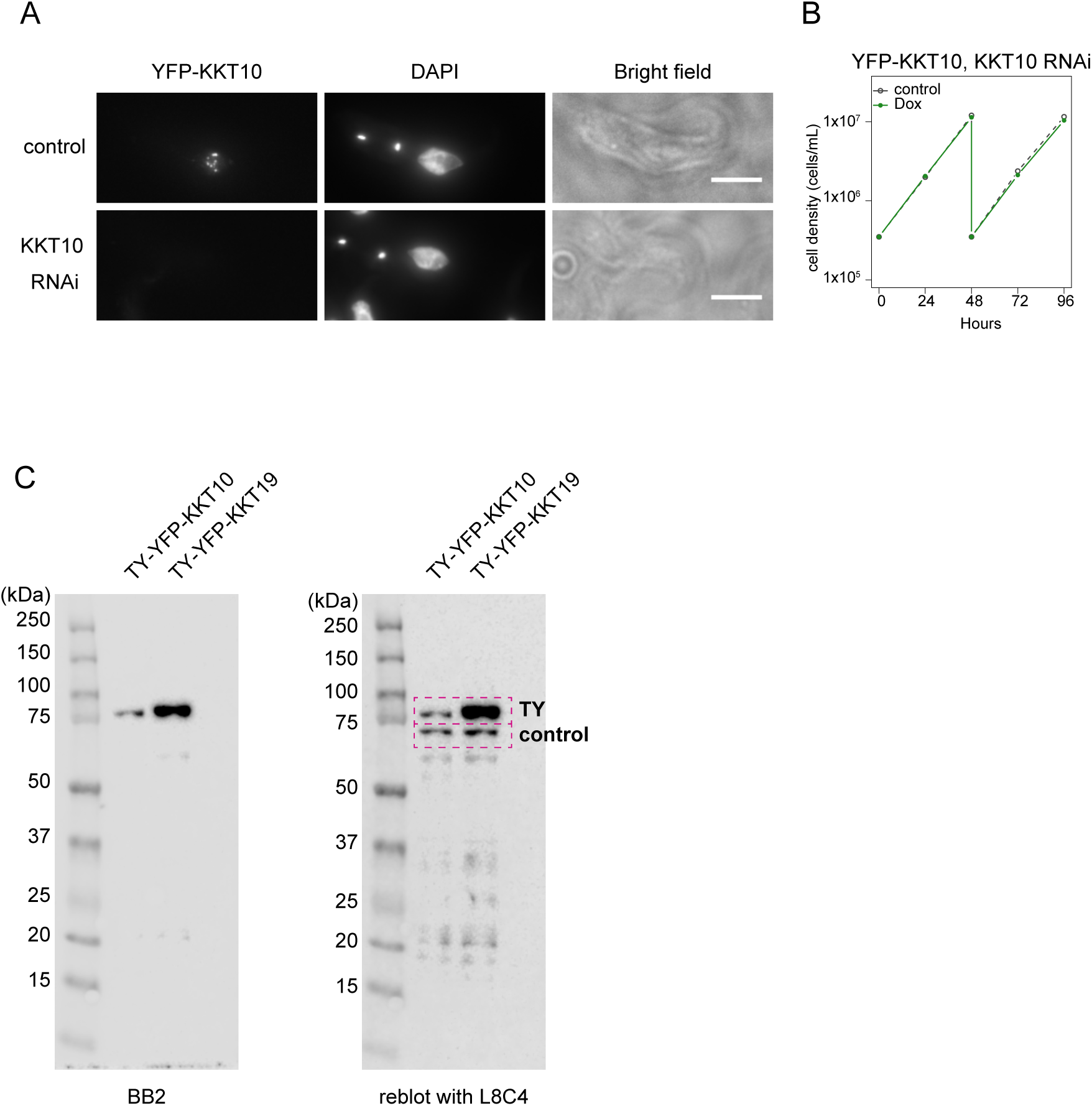
KKT10-specific RNAi does not affect cell growth. (A) Example of 2K1N cells expressing YFP-KKT10 under KKT10-specific RNAi. Cells were fixed at 24 hours postinduction of RNAi. Control is an uninduced cell culture. Maximum intensity projections are shown. Bars, 5 µm. (B) Growth curve of YFP-KKT10 with KKT10-specific RNAi. Control is an uninduced cell culture. (C) Top-bottom gels of cropped immunoblots shown in Fig. 1C.

**Figure S2.**
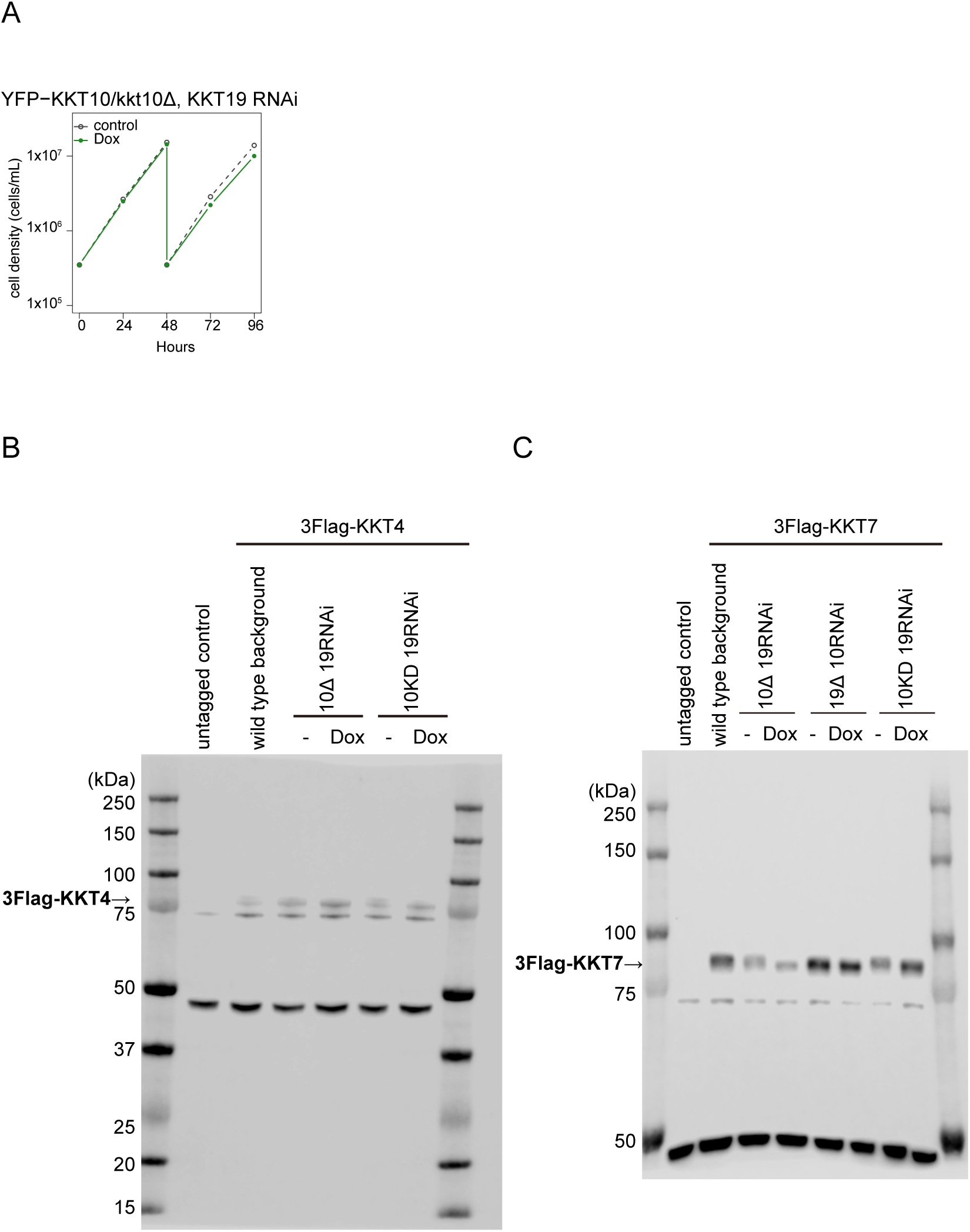
YFP-KKT10^WT^ is fully functional. (A) Growth curve of YFP-KKT10/kkt10Δ KKT19 RNAi. Control is an uninduced cell culture. Similar results were obtained from three independent experiments. (B and C) Top-bottom gels of cropped immunoblots shown in Fig. 2D. SmOxP9 was used as an untagged control.

**Figure S3.**
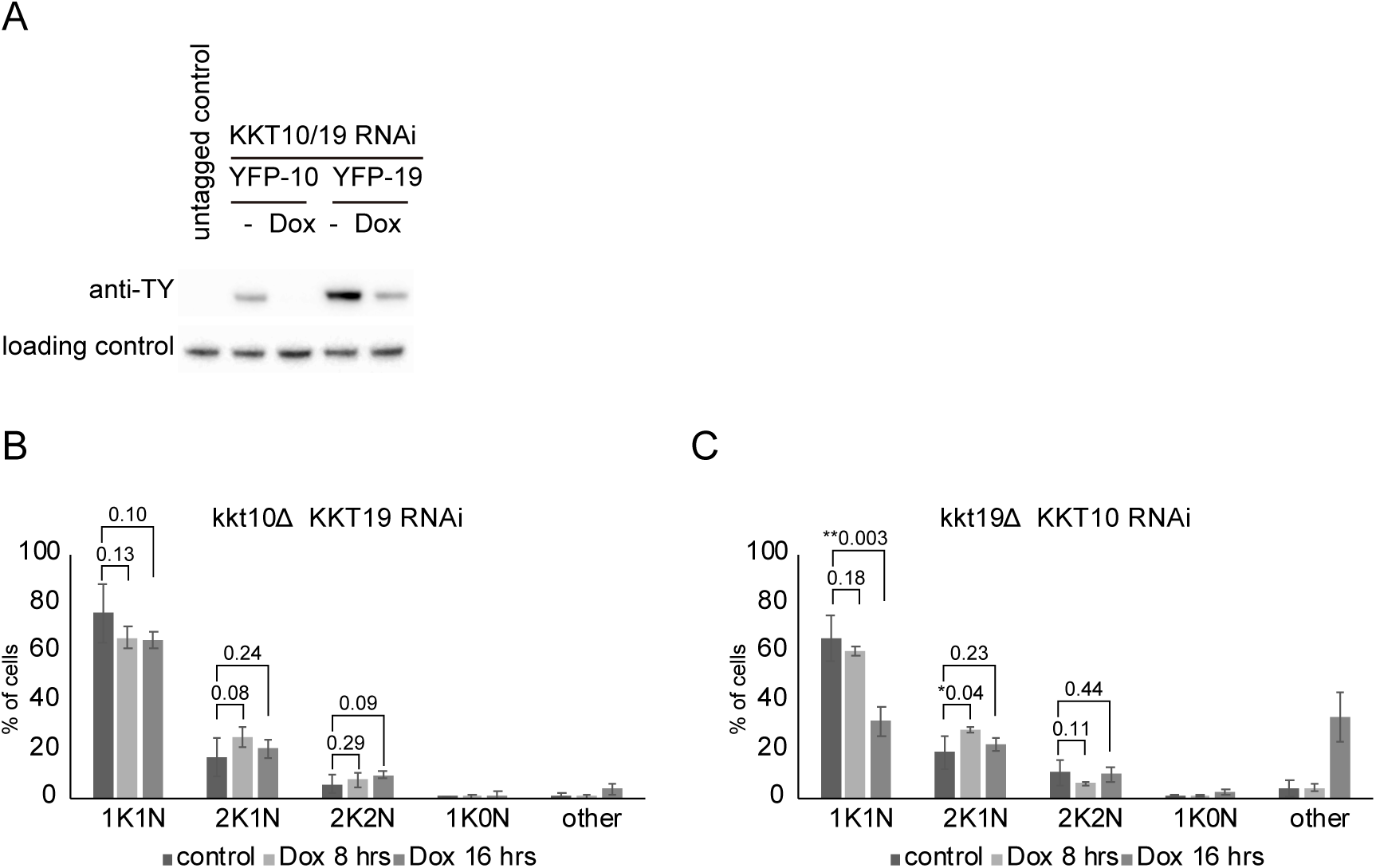
Cell cycle profiles of KKT10/19 depletion. (A) TY-YFP-tagged KKT10 and KKT19 after KKT10/19 double knockdown RNAi for 24 hours were detected by immunoblotting against the TY tag using BB2 antibodies. PFR2 detected by L8C4 antibodies was used as a loading control. SmOxP9 was used as an untagged control. (B and C) Quantification of cells with indicated DNA contents. Control is an uninduced cell culture. Error bars represent standard deviation from three independent experiments (n ≥ 314).

**Figure S4.**
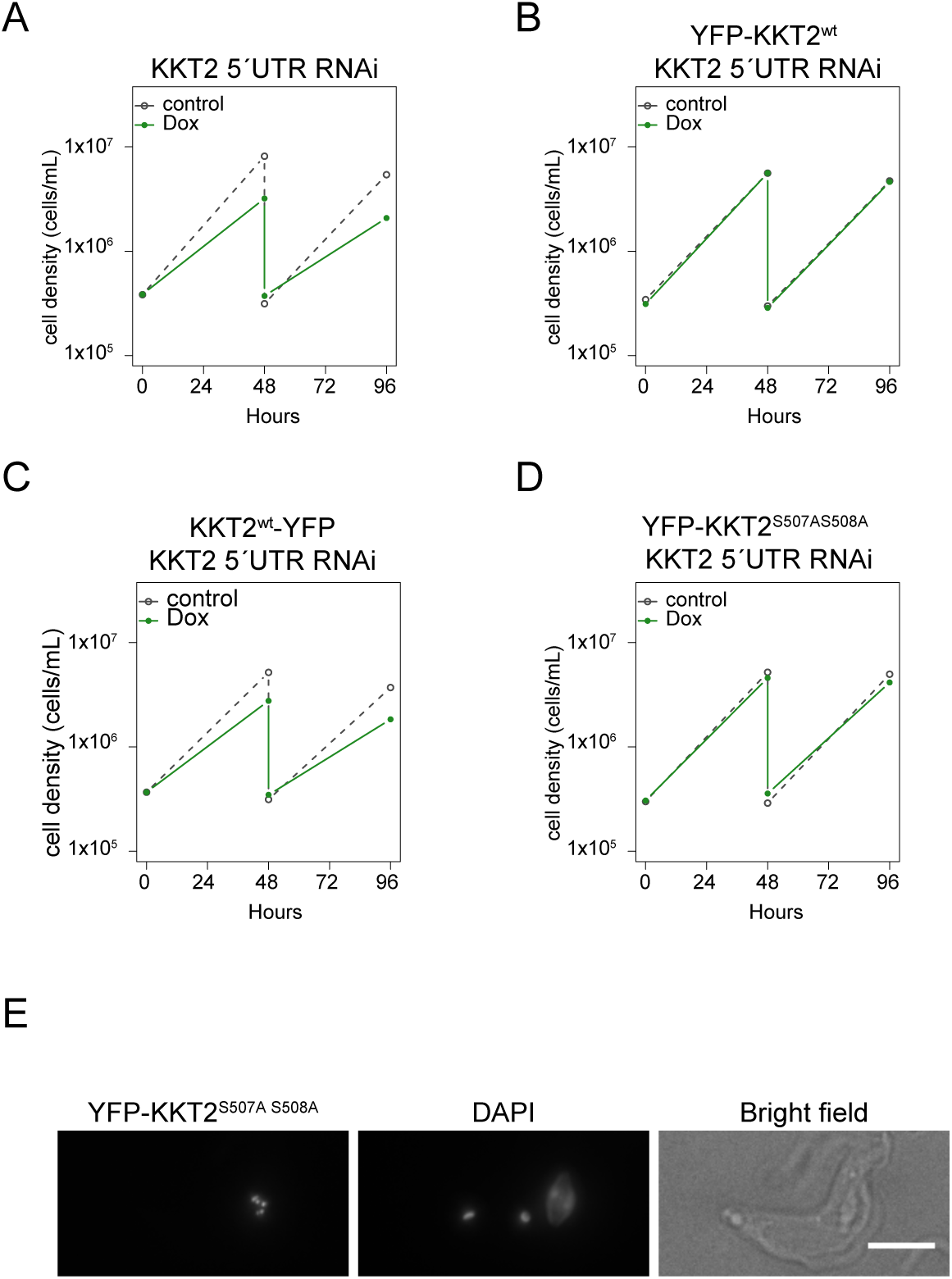
YFP-KKT2^S507A S508A^ is functional in procyclic cells. (A–D) Growth curve of (A) KKT2 5’UTR RNAi, (B) YFP-KKT2^wt^, KKT2 5’UTR RNAi, (C) KKT2^wt^-YFP, KKT2 5’UTR RNAi, and (D) YFP-KKT2^S507AS508A^, KKT2 5’UTR RNAi. Control is an uninduced cell culture. (E) YFP-KKT2^S507A S508A^ localizes normally at kinetochore. Example of cells expressing YFP-KKT2^S507A S508A^.

Table S1. Lists of trypanosome cell lines, plasmids, oligos and sequence of synthetic DNA used in this study.

Table S2. Lists of proteins identified in the immunoprecipitates of YFP-tagged KKT7N and KKT7C by mass spectrometry, Related to Figure 5.

